# Three species of *Melaleuca* from Western Australia are highly susceptible to *Austropuccinia psidii* in controlled inoculations

**DOI:** 10.1101/2022.07.25.500947

**Authors:** Alyssa M Martino, Robert F Park, Peri A Tobias

## Abstract

*Austropuccinia psidii*, the fungus causing myrtle rust, was detected in Western Australia for the first time in June 2022. Few Western Australian plant species have been screened for response to the pathogen. *Melaleuca thyoides, Melaleuca marginata* and *Melaleuca leucadendra* grown from seeds sourced from Western Australian populations were all highly susceptible to an isolate of the pathogen from eastern Australia.

---

*Austropuccinia psidii (G. Winter) Beenken comb. nov*., formerly *Puccinia psidii* (Beenken 2017) is the causal agent of myrtle rust and has been present in Australia since 2010 (Carnegie et al. 2010). Causing disease on species within the family Myrtaceae, it is present in New South Wales and Queensland (Carnegie et al. 2010; Carnegie and Lidbetter 2012; Pegg et al. 2014; Carnegie and Pegg 2018), Victoria (Agriculture Victoria 2022), Tasmania (Department of Natural Resources and Environment Tasmania 2020), the Northern Territory (Westaway 2016), and was most recently detected in Northern Western Australia (Agriculture and Food 2022).

Since the incursion into Australia, *A. psidii* has already caused the decline of keystone Myrtaceae including *Melaleuca quinquenervia* (Cav.) S.T.Blake and several rainforest understory species such as *Rhodamnia rubescens* (Benth.) Miq. and *Rhodomyrtus psidioides* (G.Don) Benth. (Carnegie et al. 2010; Carnegie and Lidbetter 2012; Pegg et al. 2014). Natural infections and pathogenicity testing has revealed over 345 susceptible species in Australia (Carnegie and Pegg 2018). Of particular concern is the susceptibility of species within Myrtaceae rich biodiversity hotspots such as that of South-West Western Australia, where myrtle rust has not yet been detected (Myers et al., 2000; Beard et al., 2000).

*Melaleuca* species are widely distributed throughout Western Australia, providing a range of ecosystem functions (Brophy et al. 2013). Previous screening of *Melaleuca* species against *A. psidii* has shown a variability in susceptibility to the pathogen (Pegg et al. 2018). As few Western Australian *Melaleuca* species have been tested for their response to *A. psidii*, the threat to these species remains largely unknown. With the recent detection of the pathogen in the Kimberley region of Western Australia on two *Melaleuca* species, yet to be formally identified (Agriculture and Food 2022), there is an urgent need to determine the vulnerability of species from Western Australia to aid conservation efforts.

To determine susceptibility of *Melaleuca* species from Western Australia in response to the pathogen, seed from *Melaleuca thyoides* Turcz. and *Melaleuca marginata* (Sond.) Hislop, Lepschi & Craven was obtained from Department of Biodiversity, Conservation and Attractions King’s Park seed bank. Although known to be susceptible (Pegg et al. 2018), seed from and *Melaleuca leucadendra* (L.) (L.) was also obtained as the isolated population not been assessed for susceptibility. For each species, seed was collected from multiple trees at 33°39’45”S 115°20’59”E, 29°07’21.9”S 115°05’46.8”E, and 16°47’10.0”S 124°55’14.5”E respectively (Figure 1).

**Figure 1.**
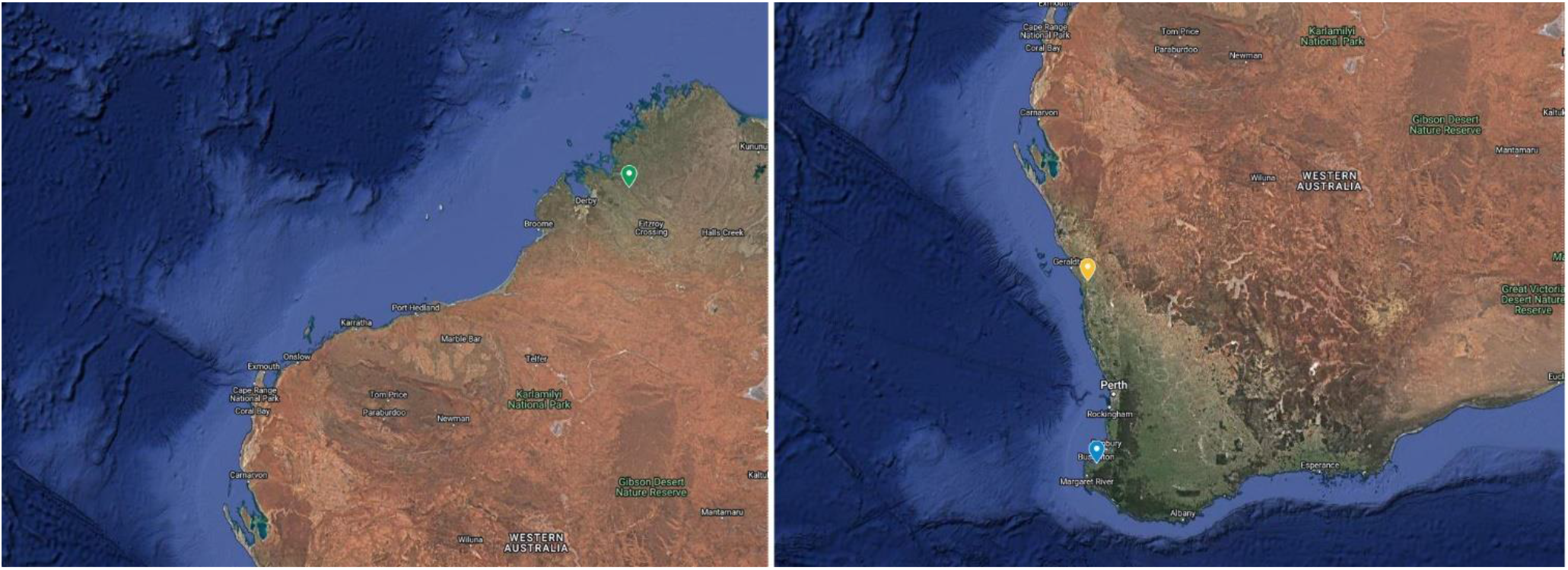
Seed collection sites for *Melaleuca leucadendra* (green tag), *Melaleuca thyoides* (yellow tag), and *Melaleuca marginata* (blue tag). Seed was collected from multiple parents at each site. Image generated in Google My Maps.

Seeds were sown into perforated trays containing a mix of 2:1:1 peat, coconut coir, and perlite supplemented with Osmocote® Native Controlled Release Fertiliser then covered with a fine coating of vermiculite. Perforated trays were placed into solid trays filled with and always maintaining 1 cm of water. Seeds were germinated under natural light in a climate-controlled glasshouse set at 26°C/20°C daytime/night time temperature on a 12 hour cycle. Germinated seedlings were transplanted into 85 mL pots (5 cm diameter and depth) containing a mix of 2:1:1 Osmocote® Native Premium Potting Mix, peat, and perlite supplemented with Osmocote® Native Controlled Release Fertiliser then placed in solid trays filled with and always maintaining 1 cm of water. Seedlings were grown under the same glasshouse conditions as for germination.

Inoculation of 57 *M. thyoides*, 26 *M. marginata* and 51 *M. leucadendra* was conducted three months post germination at the Plant Breeding Institute at the University of Sydney (Cobbitty, NSW) alongside a known highly susceptible host *Syzygium jambos* as a positive control. Approximately 50 mg of *A. psidii* urediniospores from a single isolate (Sandhu and Park 2013) was added to 50 mL of Isopar® for a final concentration of 1 mg spores/mL. Seedlings were inoculated with inoculum using an aerosol sprayer and relocated to a humid incubation chamber for 24 hours at 20°C. After incubation, seedlings were transferred to a glasshouse with the temperature set to 26°C/20°C daytime/nighttime temperature on a 12 hour cycle.

Within 12 days post inoculation, symptoms had developed on highly susceptible plants. Plants were scored using the system developed by Morin et al. (2012), ranging from completely resistant (score 1) to highly susceptible (score 5). For all species, spores appeared on leaves, stems, and petioles in highly susceptible plants (Figure 2). The highest level of susceptibility was recorded in *M. marginata* with 76.9% of plants highly susceptible and 23.1% showing no symptoms (Table 1). While the majority of *M. leucadendra* assessed were highly susceptible (62.8%), some plants showed no signs of infection (29.4%) while others showed variable levels of response to the pathogen (Table 1). 35.1% of *M. thyoides* assessed were highly susceptible with 45.6% showing no symptoms and others showing variable response to the pathogen (Table 1).

**Table 1.**
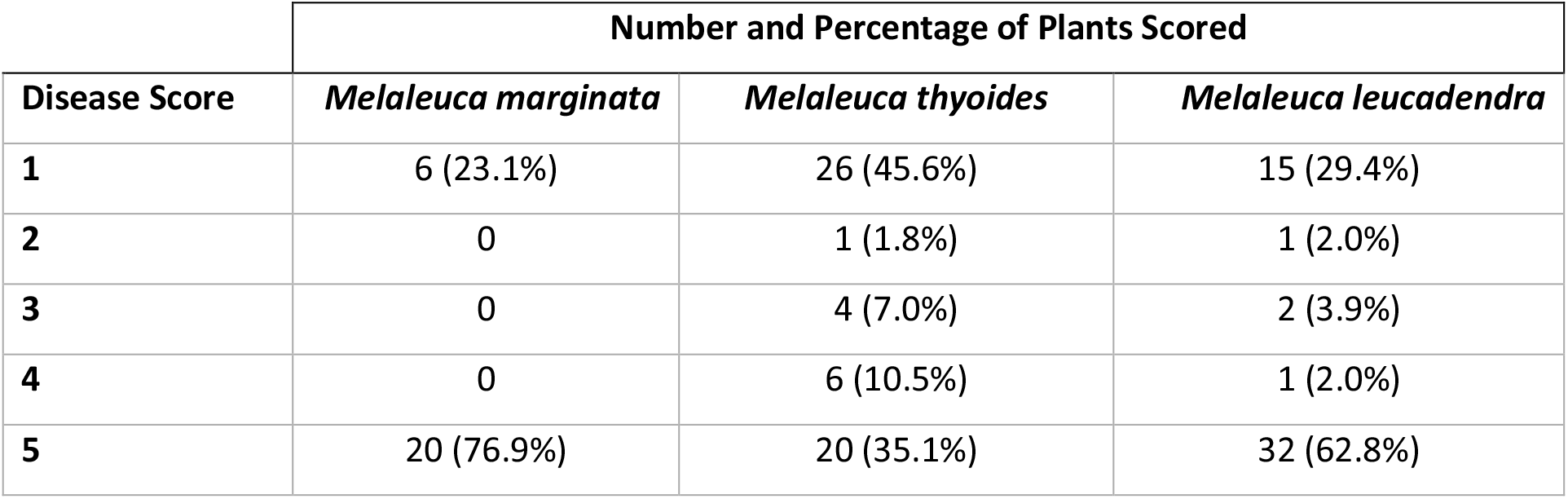
Disease scoring, based on Morin et al. (2012), of controlled inoculation of *Austropuccinia psidii* of *Melaleuca thyoides, Melaleuca marginata, Melaleuca leucadendra* and the percentage of plants observed in each disease scoring category.

| Disease Score | Number and Percentage of Plants Scored |  |  |
| --- | --- | --- | --- |
|  | *Melaleuca marginata* | *Melaleuca thyoides* | *Melaleuca leucadendra* |
| 1 | 6 (23.1%) | 26 (45.6%) | 15 (29.4%) |
| 2 | 0 | 1 (1.8%) | 1 (2.0%) |
| 3 | 0 | 4 (7.0%) | 2 (3.9%) |
| 4 | 0 | 6 (10.5%) | 1 (2.0%) |
| 5 | 20 (76.9%) | 20 (35.1%) | 32 (62.8%) |

**Figure 2.**
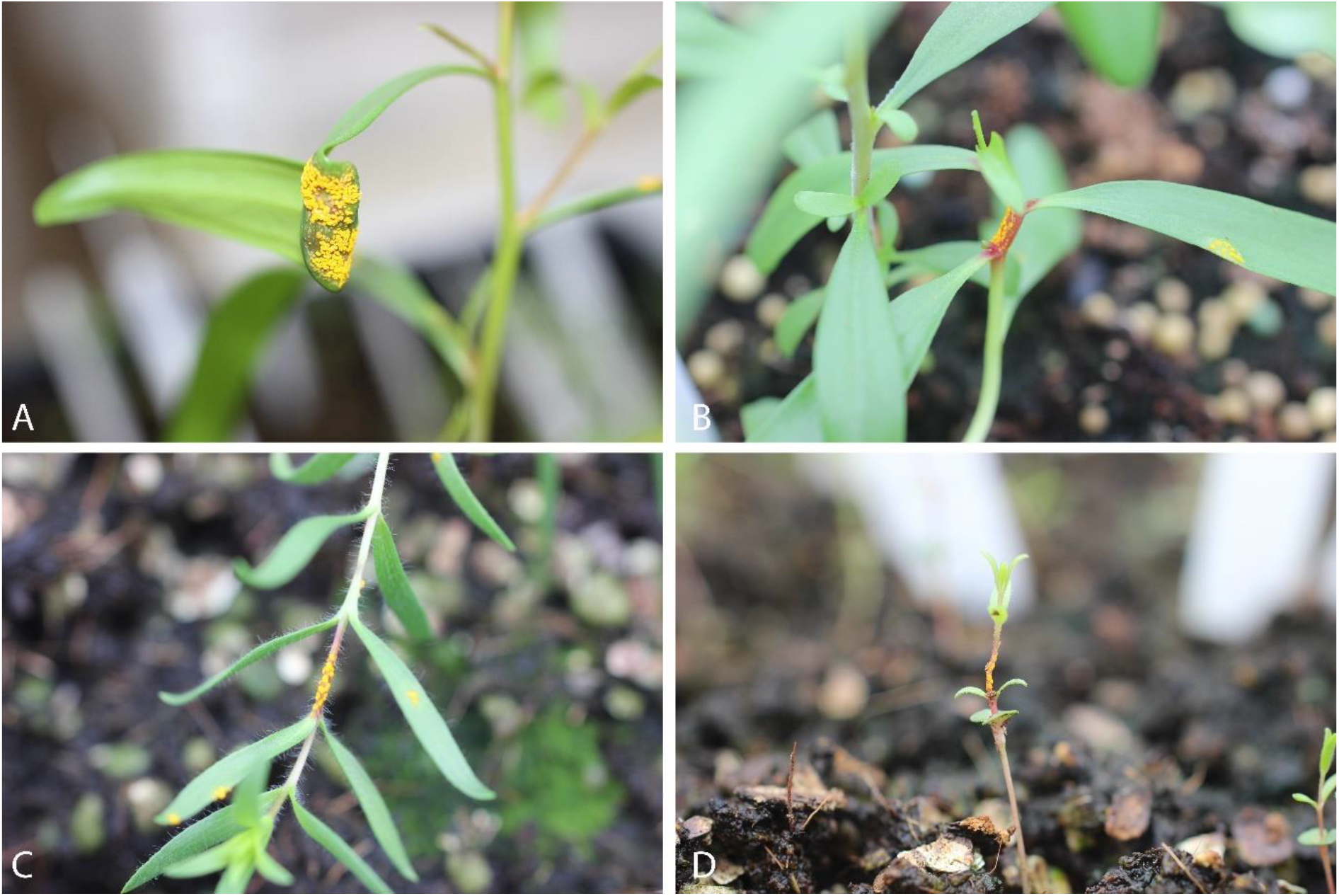
Disease symptoms of controlled inoculation of *Austropuccinia psidii* on (A, B) *Melaleuca leucadendra*, (C) *Melaleuca marginata*, and (D) *Melaleuca thyoides* at 14 days post inoculation.

These results indicate a high level of susceptibility in all three species to *A. psidii*, and for the first time reveal susceptibility in *M. thyoides* and *M. marginata*. With the pathogen now present within Western Australia, these results highlight the need to monitor myrtaceous species in the native ecosystem. The results also underscore the importance of screening more Western Australian species to determine their vulnerability to the pathogen to aid conservation efforts.

## Acknowledgements

This work was funded by the Australian Research Council under linkage project LP190100093 and The University of Sydney. Thank you to the Department of Biodiversity, Conservation and Attractions (King’s Park) for suppling the seed for this work. Thank you to Bob Makinson for the introductions that facilitated this work going ahead.

